# Damage and Misrepair Signatures: Compact Representations of Pan-cancer Mutational Processes

**DOI:** 10.1101/2025.05.29.656360

**Authors:** Caitlin F. Harrigan, Kieran R. Campbell, Quaid Morris, Tyler Funnell

**Affiliations:** Department of Computer Science, University of Toronto, Toronto, Canada; Vector Institute, Toronto, Canada; Lunenfeld-Tanenbaum Research Institute, Toronto, Canada; Memorial Sloan Kettering Cancer Center, New York, USA; Department of Hematology and Hematopoietic Cell Transplantation, City of Hope National Medical Center, Los Angeles, USA

**Keywords:** Mutational signatures, Probabilistic modelling, Cancer evolution

## Abstract

Mutational signatures of single-base substitutions (SBSs) characterize somatic mutation processes which contribute to cancer development and progression. However, current mutational signatures do not distinguish the two independent steps that generate SBSs: the initial DNA damage followed by erroneous repair. To address this modelling gap we developed DAMUTA, a hierarchical Bayesian probabilistic model that infers separate signatures for each process, and captures their sample-specific interaction. We applied DAMUTA to 18,974 pan-cancer whole genome sequencing mutation catalogues from 23 cancer types and show that tissue-specificity in mutation patterns is driven largely by variability in damage processes. We also show that misrepair processes are predictive of DNA damage response deficiencies. Unlike existing approaches, DAMUTA distinguishes damage from misrepair contributions, and we demonstrate significant improvements over a mutational-burden baseline or signatures from the COSMIC database. Our analysis reveals a shared pan-cancer pattern of early clonal transition-mutations which shifts to a more uniform substitution pattern consistent with increased reliance on translesion synthesis for damage tolerance. DAMUTA thus generates a compact set of signatures which resolves redundancies of current signature models, disentangles the effects of DNA damage and misrepair processes, and facilitates improved stratification of tumours, while providing a framework towards a unified pan-cancer model of the cellular response to DNA damage.

## Main

Somatic mutations accumulate throughout tumour development as a result of exogenous and endogenous mutational processes; these processes can result in distinct patterns of mutations called “signatures” ^1,19,31^. Most somatic mutations are single base substitutions (SBS) and hundreds of SBS signatures have been described ^1,8^. Some signatures associated with underlying DNA repair deficiencies predict patient prognoses or therapy sensitivities ^4^ and their activity can help guide treatment. For example, HRDetect uses signature activities to identify homologous recombination-directed repair (HR) deficient tumours; these tumours respond well to PARP inhibitors and platinum-based chemotherapy ^5,7,20,21^.

The Catalogue of Somatic Mutations in Cancer (COSMIC) ^30^ associates SBS signatures with etiologies that have experimental or epidemiological support. Some are well-characterized such as the C-to-G and C-to-T substitutions in a YTCA context associated with activity of the APOBEC3 family of cytosine deaminases ^10^ and their corresponding SBS signatures, SBS2 and SBS13 which are highly active in many cancers ^1,23^. However, most SBS signatures lack clear etiologies ^1^ and few COSMIC signatures have been definitively linked to loss-of-function of particular repair proteins. One challenge in making these linkages is that current SBS signatures model the combined result of damage and misrepair, thereby implicitly assuming a one-to-one mapping between damage and misrepair processes, whereas there is substantial redundancy in DNA repair systems ^14,34^. Thus, a DNA repair deficiency in the context of different types of damage will result in a different SBS signature ^36^. It is therefore difficult for SBS signature analyses to isolate the contributions of individual damage or misrepair process activities ^13,17,24^. Failure to capture these individual contributions may be the cause of tissue-specific manifestations of core COSMIC signatures ^8^, and reported differences between the patient-derived SBS signatures and ones derived from experimental perturbations to DNA repair pathways ^35,39^. Here we propose a more expressive and interpretable mutation signature model that captures the individual contributions of damage and misrepair to mutation rates, thereby permitting tissue- and sample-specific inference of the interactions between these processes.

This new signature model, DAMUTA (Dirichlet Allocation of Mutations), defines distinct damage and misrepair signatures, and models mutation type frequencies as the result of interactions between these two types of signatures. By representing the two-step process of mutation generation (Fig. 1a,b), DAMUTA is able to capture damage or misrepair-specific patterns. Similar efforts to separate the effects of damage and defective repair include RepairSig, which estimates “secondary” mutational signatures associated with repair defects ^37^. RepairSigs assumes fixed and known primary damage signatures, whereas DAMUTA estimates signatures *de novo* and imposes additional constraints on misrepair signature profiles to overcome challenges in identifiability when fitting two sets of signatures ^37^.

**Figure 1.**
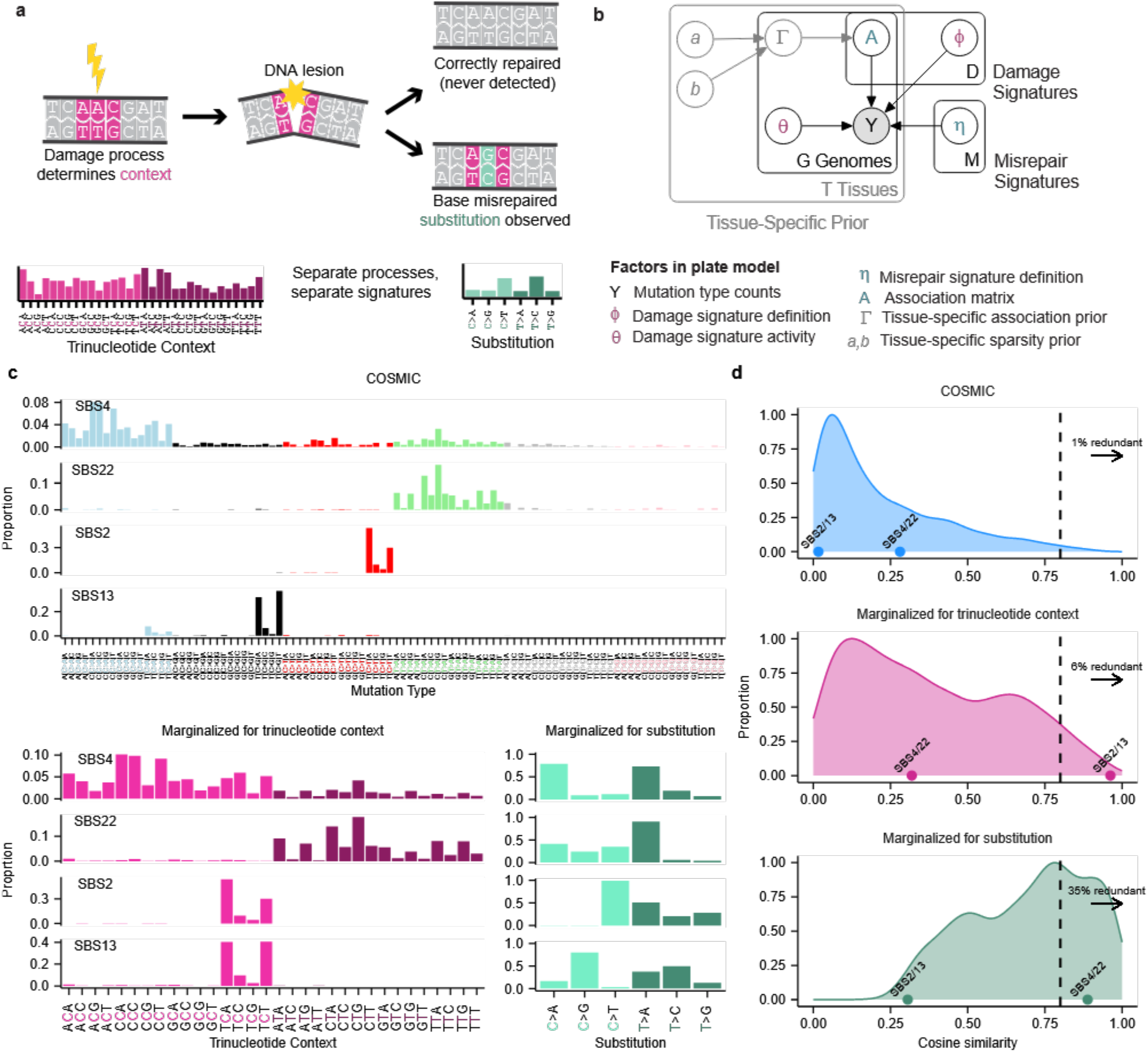
Method overview. **a,** Single base mutations are generated by two sequential events: the DNA damage process selects a sequence context where a lesion occurs, then the misrepair process results in a substitution in this context. **b**, DAMUTA probabilistic graphical model. Parameters related to Damage signatures are indicated in maroon, parameters related to Misrepair signatures are indicated in green. An optional hierarchical tissue-specific prior is indicated in grey. **c**, Examples of four COSMIC signatures marginalized to trinucleotide context and substitution profiles. **d**, Density plots of cosine similarity between n = 1770 pairs of 60 full and marginalized, non-artifact COSMIC SBS signatures. Signature pairs which have >0.8 cosine similarity are deemed “redundant”, the proportion of which is annotated.

To illustrate the benefits of separately modelling damage and misrepair, we used DAMUTA to distinguish perturbations to APOBEC3-mediated damage and modulation of UNG-initiated base excision repair in breast and B-cell lymphoma cell lines, whereas these perturbations could not be distinguished using COSMIC SBS activities.

We also show that DAMUTA leads to a more compact and useful representation of other mutational processes. Because it removes the redundancy in standard SBS signatures, DAMUTA only requires 24 signatures to capture variation in a pan-cancer cohort, comparing to 78 COSMIC SBS signatures. We show that despite its smaller feature set, classifiers trained on DAMUTA’s 24 signature activities were consistently better at detecting alterations in DNA repair pathways than ones trained on the 78 COSMIC signature activities.

We provide DAMUTA signatures and activities for 18,974 whole genome sequenced tumours originating from 23 organs (Fig. S1), and describe their activity distributions across tissue types and cancer evolution. We also describe common pan-cancer damage-misrepair connection patterns, and use these patterns to reveal how damage and misrepair may interact to generate the reported COSMIC SBS signatures. Together, our study demonstrates the utility of modelling somatic mutations as the consequence of interacting sequential processes, as a means to discover and disentangle the distinct effects of DNA injury and misrepair, via a more compact representation of mutational signatures in cancer.

## Results

### Conventional SBS signatures are composed of redundant sequence context and substitution profiles

Analysis of COSMIC signatures showed patterns of redundancy in their representation of either damage, via their trinucleotide contexts, or misrepair, via their substitution profiles. For example, SBS4 and SBS22 have similar substitution profiles but differ in their trinucleotide context, while SBS2 and SBS13 have similar trinucleotide contexts but different substitutions (Fig. 1c). To quantify this redundancy we computed trinucleotide and substitution profiles through marginalization for 30 COSMIC v2 and 60 of the 78 COSMIC v3.2 SBS signatures that represent non-artifact signatures (*e.g.* Fig. 1c *bottom*), and measured pairwise cosine similarities between the full SBS signatures, their trinucleotide contexts, or their substitution types. We labelled signatures as “redundant” if they shared >0.8 cosine similarity. COSMIC v3.2 signatures had low redundancy, 1%, (Fig. 1d *top*, Supplementary Fig. S2, median cosine similarity=0.14) reflecting an effort to reduce their redundancy from COSMIC v2^1^ which had 3% redundancy (median cosine similarity=0.26).

Despite the low overall signature redundancy, we found 6% of COSMIC v3.2 (hereafter COSMIC) trinucleotide context profiles were redundant (Fig. 1d *middle*, median cosine similarity=0.33) including SBS2 and SBS13, and 35% of substitution profiles were redundant (Fig. 1d *bottom*, median cosine similarity=0.73) including SBS4 and SBS22. Neither of these pairs of signatures were redundant in their COSMIC-SBS form. This partial redundancy within COSMIC SBS signatures suggests they could be more compactly represented as specific combinations of trinucleotide context and substitution profiles, and that there are likely fewer distinct substitution profiles than sequence context profiles.

### DAMUTA models mutational signatures as combinations of damage and repair processes

DAMUTA is a hierarchical Bayesian model that separately represents damage and misrepair processes, and models how their interactions lead to mutations in order to explicitly describe the two-step nature of mutational processes (Fig. 1b). DAMUTA defines the damage process as the factor that selects the trinucleotide context of DNA damage, and the misrepair process as the one that selects the mutation by which the damage is resolved. DAMUTA thus defines two sets of signatures: Damage signatures, *ϕ,* over 32 trinucleotide contexts centred on the pyrimidine-normalized alteration (*i.e.* ACA, ACC, …, TTT); and Misrepair signatures, *η*, over 6 categories representing the substitution of the pyrimidine, *i.e.*, C>A, C>G, C>T, T>A, T>C, or T>G. Hereafter, capitalized “Damage” and “Misrepair” refer to signatures estimated by DAMUTA.

DAMUTA captures the relationship between Damage signature *ϕ*_*d*_ and Misrepair signature *η*_*m*_ for genome *g* in the entry *A*_*g,d,m*_ of the association matrix *A*_*g*_ (see Fig. 1b), each row of which is Dirichlet distributed. In order to ensure identifiability of the model, we introduce regularization of *A* via a tissue-specific sparsity prior Γ (Fig. 1b), with potentially tissue-specific parameters a and b. When the tissue type is unknown, we default to an uninformative prior for the rows of A (Supplementary Fig. S3).

Damage and Misrepair signatures are global latent variables, the Damage signature activities *θ*_*g*_ and association matrix probabilities *A*_*g*_ are sample-specific to capture the tissue-, cell-, and sample-specific activity and interaction of damage and misrepair processes.

To analyse the interaction of damage and misrepair processes, we define an *interaction matrix W_g_* = *θ*_*g*_*A*_*g*_, *i.e.,* the association matrix for the sample whose columns are scaled by the activity of their associated damage signature. Misrepair signature activities are computed by taking the row sums of *W*_*g*_.

### 18 Damage Signatures & 6 Misrepair Signatures summarise the SBS profiles of 18,947 tumours

We fit DAMUTA *de-novo* to 442,701,158 somatic single base substitutions from 18,947 whole genome sequenced primary and metastatic tumours, finding that a model with 18 Damage (Fig. 2a) and 6 Misrepair signatures (Fig. 2b) optimized data fit and consistency according to standard model selection criteria (Methods, Supplementary Figs. S4, S5). Damage signatures tended to have higher weight in either NCN or NTN contexts, and not both: 12/18 are C-context biased, 5/18 are T-biased, and 1/18 is C/T balanced. A similar bias is found in COSMIC signatures: 32/60 are C-biased, 19/60 are T-biased, and 9/60 are C/T balanced (Supplementary Fig. S6).

**Figure 2.**
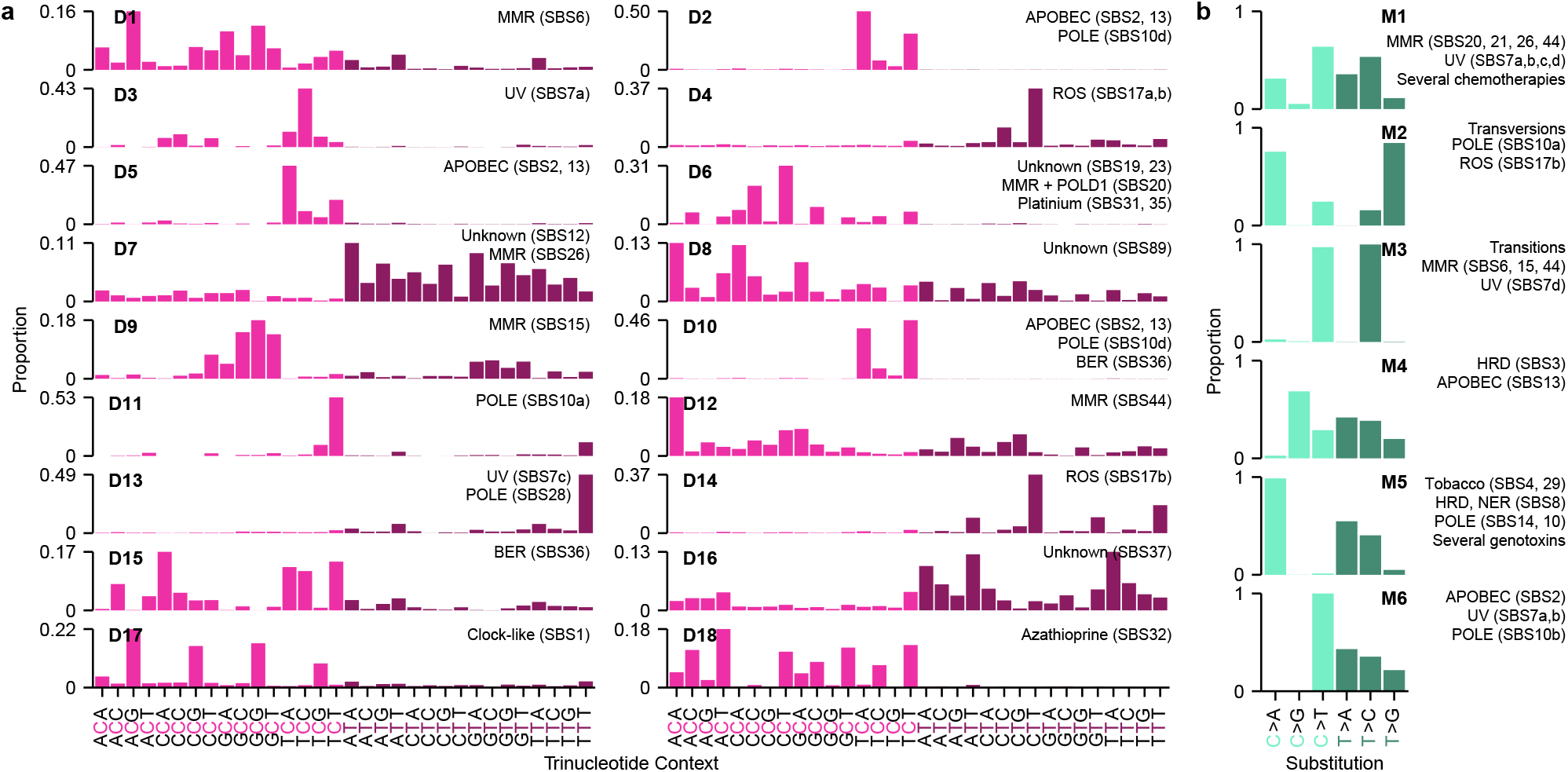
Composition of 18 DAMUTA Damage signatures and 6 Misrepair signatures derived from 18,947 tumours. **a**, Damage signatures with proposed aetiologies annotated from COSMIC signatures with ≥ 0.85 cosine similarity. MMR: Mismatch repair deficiency, BER: Base excision repair deficiency, POLE: Polymerase epsilon deficiency, POLD1: Polymerase delta deficiency, APOBEC: Apolipoprotein B mRNA editing catalytic polypeptide-like protein, UV: Ultraviolet radiation, ROS: Reactive oxygen species. **b**, Misrepair signatures with proposed aetiologies annotated from COSMIC signatures with ≥ 0.85 cosine similarity.

We labelled Damage and Misrepair signature aetiologies based on COSMIC signatures by matching the trinucleotide and substitution profiles with the highest cosine similarity. Often, as highlighted in Fig. 1c, multiple COSMIC profiles had high similarity to the same *de-novo* extracted DAMUTA signature, and we list aetiologies corresponding to COSMIC signatures with at least 0.85 cosine similarity. With the catalogue of DAMUTA signatures extracted from this pan-cancer data, we proceeded to investigate the utility of DAMUTA signature activities to investigate mutational processes.

### DAMUTA identifies Damage and Misrepair activity variation associated with *APOBEC3* and *UNG* perturbation

We tested DAMUTA using *APOBEC3* knockout and *UNG* modulation data from Petljak *et al* ^23^. The standard model for APOBEC signature generation involves exposed single stranded DNA acting as substrate for APOBEC3, which deaminates cytosine to uracil (Supplementary Fig. S7). The edited base is most often misrepaired to thymine, thus introducing C>T substitutions. Uracil-DNA glycosylase (UNG) is a base excision repair (BER) protein known for its role in resolving APOBEC-introduced abasic sites ^26^ and is implicated in the generation of APOBEC-like mutations ^23^. Mutation counts from two breast cancer cell lines (MDA-MB-453 and BT-474) and one B-cell lymphoma cell line (BC-1) were analyzed using DAMUTA. These cells were either wildtype, *APOBEC3A* knockout, *APOBEC3B* knockout, or *APOBEC3A*/*APOBEC3B* double knockouts. Because breast cells constitutively express *UNG, UNG*-wild-type experiments in this cell type were labelled as UNG-high, and the *UNG*-knockout as UNG-low. Conversely, the B-cell lymphoma cell line naturally exhibits low *UNG* expression, and these were labeled UNG-low (wildtype) or UNG-high (after UNG-GFP viral transduction).

We fixed signature definitions to those extracted from patient tissue samples (Fig. 2), and estimated Damage and Misrepair signature activities *de novo* for individual cell line samples (Fig. 3). We found distinct patterns of Damage and Misrepair activity across experimental settings (Fig. 3a). Damage signature activities clearly separated the *APOBEC3A* knockout and *APOBEC3A*/*APOBEC3B* double knockout cell lines from the wildtype and *APOBEC3B* knockouts (Fig. 3b), in accordance with reported activities of APOBEC3A and APOBEC3B. Similarly, Misrepair signature activities completely separated samples by *UNG* status (Fig. 3c), with the wildtype B-cell lymphoma cell line, which naturally exhibits low *UNG* expression, clustered with the breast *UNG*-knockouts. This separation is not apparent when clustering by COSMIC activities, which only capture the compound effect of damage and misrepair (Supplementary Fig. S8).

**Figure 3.**
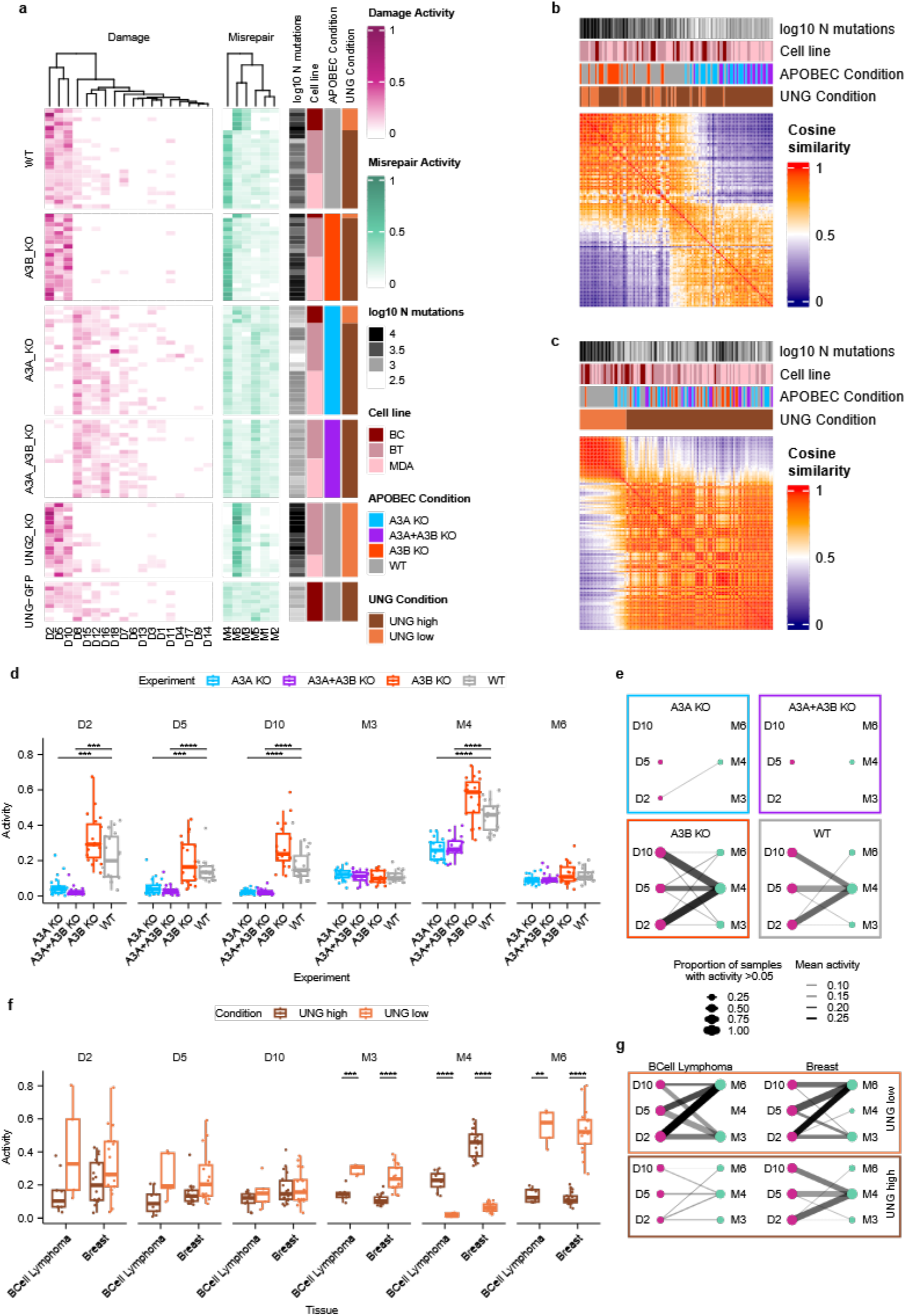
DAMUTA distinguishes perturbations to damage and misrepair processes in APOBEC mutagenesis. **a**, Activities of DAMUTA signatures in three cancer cell lines MDA: breast, BT: breast, BC: B-cell lymphoma. WT: wild-type, A3A: *APOBEC3A*, A3B: *APOBEC3B*, KO: knockout. **b**, Cosine similarities of Damage activities. Samples are ordered by PCA seriation. Annotation track colors as in **a. c**, Cosine similarities of Misrepair activities. Samples are ordered by PCA seriation. Annotation track colors as in **a. d**, Distribution of activities of six APOBEC-associated signatures in APOBEC knockout experiments in two breast cell lines. Only significant differences to wild-type are shown: *: P<0.05, **: P < 0.01, ***: P < 0.001, ****: P < 0.0001, Bonferroni-corrected, two-tailed t-tests. Boxes show interquartile range (IQR) with median line. Whiskers: 1.5 × IQR. **e**, Connection patterns between APOBEC-associated signatures across APOBEC knockout conditions. Only activities above 0.05 are visualized, for selected signatures. **f**, Distribution of activities of APOBEC-associated signatures in *UNG* knockout and reconstitution experiments in two breast cancer, and one lymphoma cell line. Boxes, whiskers, and statistical significance as in **b**. Only significant differences are shown. **g**, Connection patterns between APOBEC-associated signatures across *UNG* knockout and reconstitution conditions. Only activities above 0.05 are visualized, for selected signatures. Legend is shared with **e**.

We next focused on Damage signatures D2, D5, D10 and Misrepair signatures M3, M4, M6 for further analysis due to their similarity to COSMIC APOBEC-related signatures (Fig. 2). We observed *APOBEC3A* single and *APOBEC3A*/*APOBEC3B* double knockouts had significantly reduced activity of D2, D5, D10, M4 and M6 compared to wildtype (Bonferroni-corrected t-test, P<0.001, Fig. 3d).

In addition to examination of the Damage and Misrepair activities in isolation, the DAMUTA model supports consideration of the interactions between these processes. In Figure 3e, we visualize the connections between Damage and Misrepair captured in the interaction matrix as a bipartite graph. Although M3/M4/M6 are active in the KO samples *APOBEC3A* (Fig. 3d), their connection to D2/D5/D10 is almost entirely lost (Fig. 3e).

We next evaluated the effect of *UNG* expression on Damage and Misrepair signature activities. Overexpression of *UNG* in the B-cell lymphoma line resulted in a phenotype more similar to the wild-type breast cells, in terms of Misrepair activities (Brown UNG-high boxes, Fig. 3f), and in the connectivity observed between Misrepair and Damage signatures (Fig. 3g). This is consistent with the distinct roles of APOBEC and UNG proteins in influencing mutation patterns and repair mechanisms. Although the Damage activities are for the most part not significantly different, the Misrepair usage is changed according to UNG expression level. In UNG-low settings, for example, M3 and M6 have significantly (Bonferroni-corrected t-test P<0.01) higher activity in both the B-cell lymphoma cells and breast cells, while M4 activity is higher (Bonferroni-corrected t-test, P<0.0001) in UNG-high cells.

These results demonstrate DAMUTA’s unique ability to distinguish mutational patterns associated with loss or modulation of specific DNA damage and repair processes. While COSMIC signatures SBS2 and SBS13 appear with mixed activity, DAMUTA separates APOBEC3A-dependent damage, through Damage signature activity, from UNG-dependent repair outcomes through Misrepair signature activity.

### DAMUTA improves low-dimensional representations of SBS profiles

Given DAMUTA’s ability to distinguish DNA damage and misrepair processes, we hypothesized that they would be more informative features than COSMIC signature activities to predict changes in DNA damage response (Fig. 4). To compare the utility of DAMUTA and COSMIC activities, we assessed the performance of binary classifiers which used these activities as input representations of each tumour.

**Figure 4.**
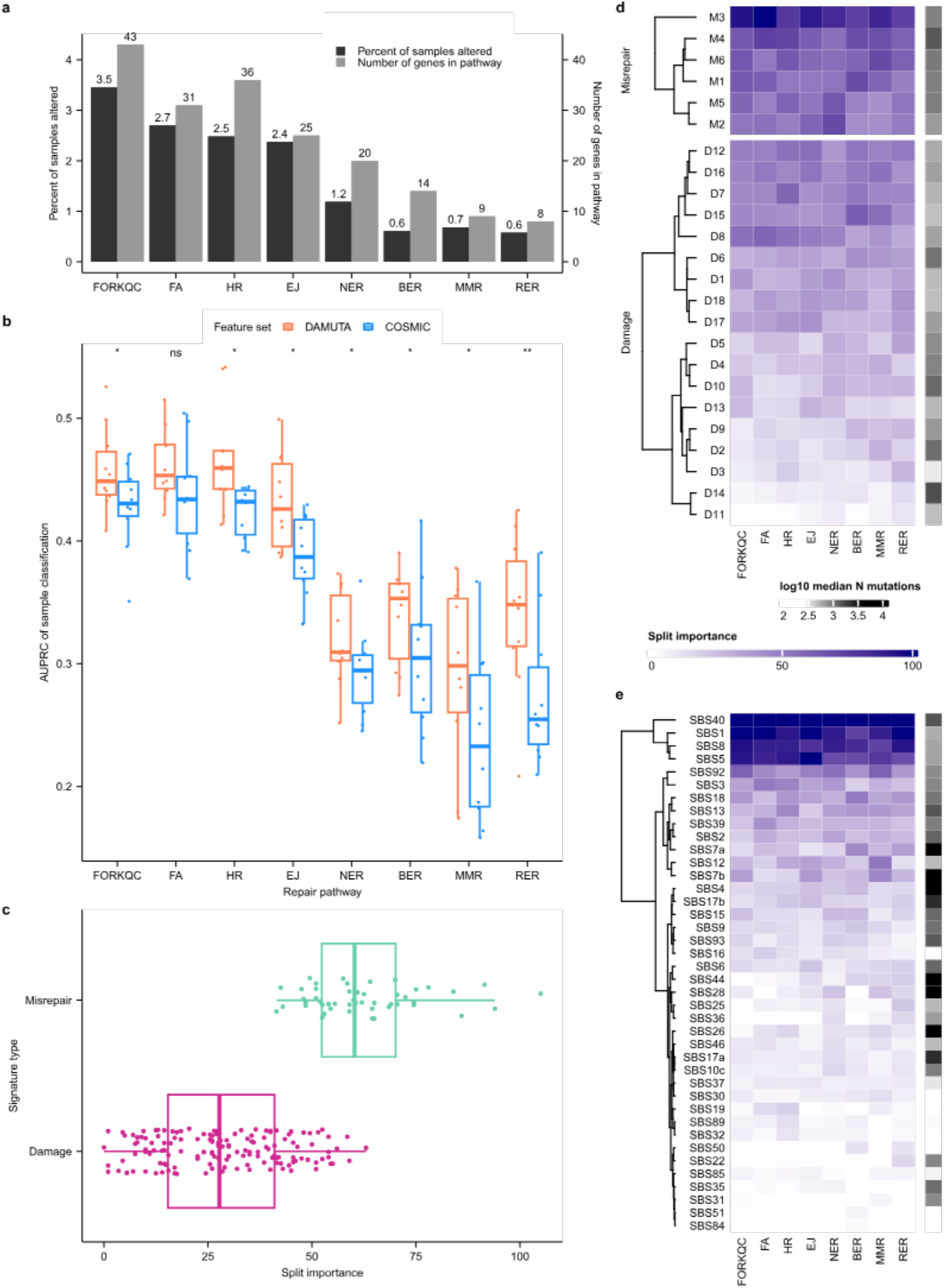
DAMUTA activities as sample classification features. **a**, Size of gene sets and percent of positive class examples for 8 DDR pathways. **b**, COSMIC vs DAMUTA feature sets performance across DDR pathways to predict non-silent point mutation in any gene in the pathway. Boxes show interquartile range (IQR) with median line. Whiskers: 1.5 × IQR. Benjamini-Hochberg adjusted paired t-test P-values indicated with ns: Not significant (P > 0.05), *: P<0.05, **: P < 0.01. Data from 10 random seeds shown. **c**, Split importances for Damage and Misrepair signatures across 8 DDR pathways. Boxes show interquartile range (IQR) with median line. Whiskers: 1.5 × IQR. **d**, Split importances of Damage and Misrepair signatures summarized for all pathways. **e**, Split importances of COSMIC signatures summarized for all pathways. Legend is shared with **d**.

Specifically, we trained gradient boosted machines (GBM) classifiers to predict per-sample alteration status of the following eight DDR pathways: DNA replication fork quality control (FORKQC), Fanconi Anemia (FA), Homologous Repair (HR), End Joining (EJ), Nucleotide Excision Repair (NER), Base Excision Repair (BER), Mismatch Repair (MMR), and Ribonucleotide Excision Repair (RER). We derived the gene lists for each of these pathways from Olivieri *et al.* ^22^, expanding the BER pathway to include NTLH1 which is implicated in the generation of SBS30^11^.

The number of genes in a pathway ranged from eight (RER) to 43 (FORKQC) (Fig. 4a). For each pathway, we marked a sample as mutated if a non-synonymous somatic point mutation was detected within any gene in that pathway. As training input, classifiers were given the estimated number of mutations assigned to either 24 DAMUTA (18 Damage, 6 Misrepair) or 78 COSMIC signatures for each of the 2,778 PCAWG ^32^ samples (see Methods). In total, we tested 16 model configurations: one for each of the eight pathways and two feature sets. Each of the 16 classifiers was re-trained 10 times, with 30% of the samples withheld for model evaluation. Data were randomly stratified on the positive class to ensure examples were evenly split across train and test sets.

We note these classification tasks were highly imbalanced: fraction of positive samples ranged from 0.6% (RER, n=16) to 3.46% (FORKQC, n=96) as shown in Fig. 4a. Therefore, we relied on the area under the precision-recall curve (AUPRC) of each classifier to evaluate performance (Fig. 4b). We found that DAMUTA features consistently out-performed COSMIC features with a mean gain in AUPRC of 13% (P < 0.001, paired t-test). In addition, the performance of the DAMUTA-based classifier was significantly different than tumor mutational burden (TMB) alone at predicting pathway mutational status (P<0.01) however this was not the case for the COSMIC-based classifier (P>0.25, Supplementary Fig. S9).

Fig. 4c compares the split feature importance values for DAMUTA Misrepair versus Damage activities, showing that on average Misrepair was twice as important as Damage, indicating that Misrepair activities indeed capture information specific to repair deficiency status (Fig. 4d). In contrast, the feature importances for COSMIC signatures were concentrated on a small number of widely active signatures (Fig. 4e, SBS1, SBS5, SBS8, SBS40), some of which broadly associated with proliferation or ageing, suggesting that the COSMIC based classifier is simply computing a surrogate for TMB.

### Recurrent Damage-Misrepair “Connection patterns” mirror some COSMIC signatures

We next sought to determine whether there were other common interaction patterns like those underlying APOBEC-associated mutations. To do so, we used non-negative matrix factorization (NMF) to decompose the 18,947 sample-specific interaction matrices into a small number of Connection patterns represented by the non-negative factors (Fig. 5a, Supplementary Figs. S10, S11). Each Connection pattern can be collapsed to a distribution over the 96 SBS types by combining Damage and Misrepair signatures using their Connection weights, and we found that some of these patterns recapitulated COSMIC signatures. Based on these comparisons, we chose to extract 30 Connection patterns, labeled C1 to C30. Increasing the number of patterns in the NMF extraction to greater than 30 did not improve their correspondence with COSMIC signatures (Supplementary Fig. S12a).

**Figure 5.**
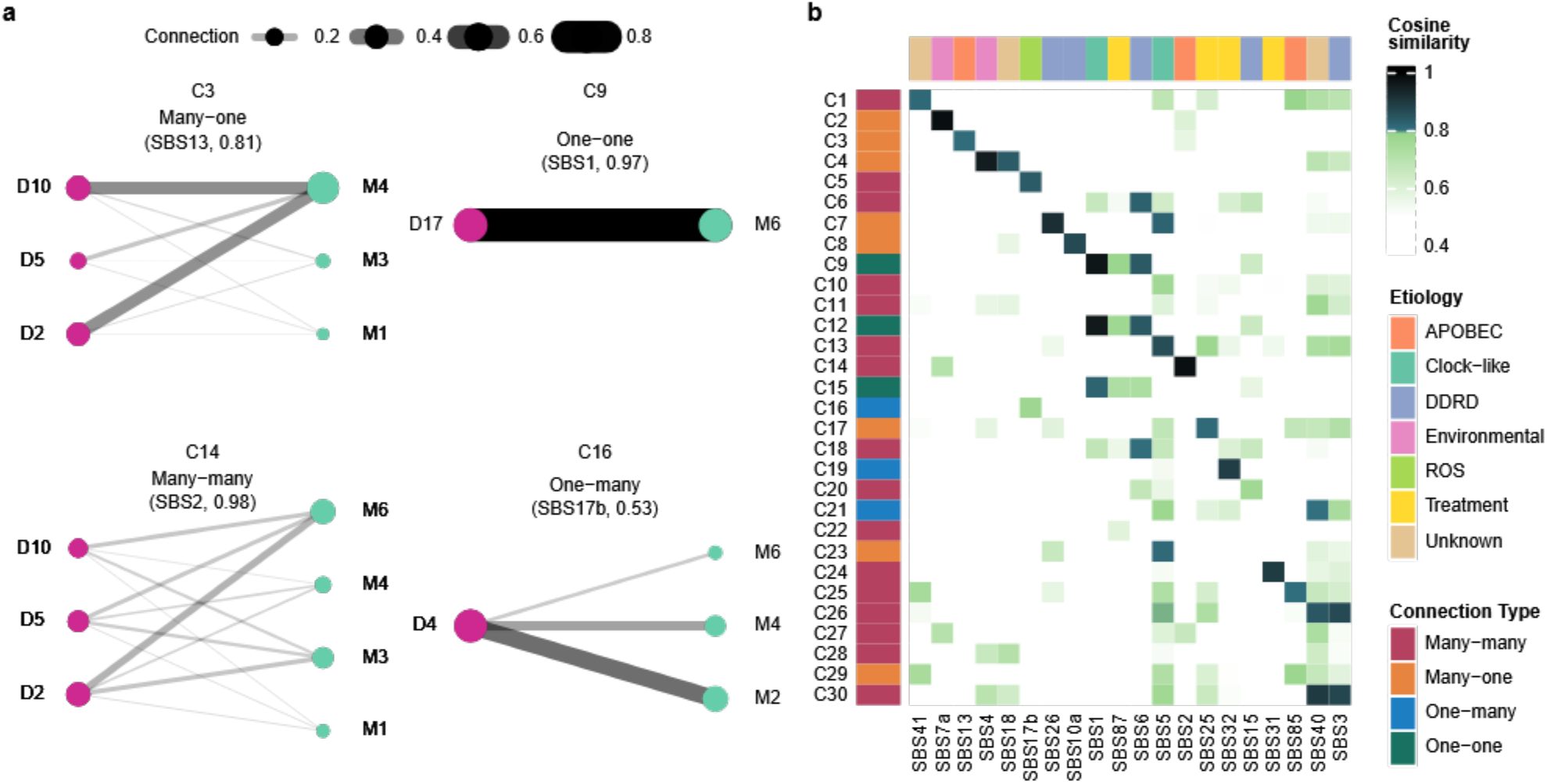
Damage-Misrepair Connection Patterns. **a**, Examples of 4 types of Connection patterns extracted using NMF. Patterns are labelled by Connection pattern name, and connection type. Only connections between Damage or Misrepair signatures with >0.05 activity are visualized. Text in parentheses indicates matched COSMIC signature and its cosine similarity based on the Connection pattern’s collapsed SBS distribution. **b**, Heatmap of cosine similarities between connection patterns and COSMIC signatures.

We next assessed the correspondence between these Connection patterns and COSMIC signatures, and matched COSMIC signatures by cosine similarity with each Connection pattern’s collapsed SBS distribution (Fig. 5b). A total of 30 COSMIC signatures had >0.8 cosine similarity to a Connection pattern (Supplementary Fig. S12b), 22 Connection patterns had >0.8 similarity to a COSMIC signature, and the remaining 8 Connection patterns were matched to their closest COSMIC signature (Supplementary Figs. S10, S12). The Damage and Misrepair signatures which we identified as relevant to APOBEC mutagenesis in cell line data (D2/D5/D10, M3/M4/M6) participated in Connection patterns C3 and C14. C3- and C14-reconstituted 96-channel SBS signatures were most similar to COSMIC APOBEC signatures SBS13 and SBS2 respectively, and in the cell lines C3 and C14 activities were positively correlated with SBS13 (R2=0.74) and SBS2 (R2=0.92) (Supplementary Fig. S13).

We next classified the Connection patterns into four types based on the number of active Damage/Misrepair signatures they connected: one-one (n=3), one-many (n=3), many-one (n=8), and many-many (n=16); Fig. 5a shows examples of each type. The high frequency of many-many, one-many, and many-one interaction types, and their correspondence with COSMIC SBS signatures, suggests that COSMIC signatures typically capture the interactions among multiple damage and/or misrepair processes.

### Damage processes are more tissue-specific than misrepair processes

Cancers arising from the same organ system or tissue type often have characteristic COSMIC mutational signature activities and evolutionary trajectories ^12,28^. To assess the tissue specificity of DAMUTA signatures, we considered both their overall activities (*i.e.*, the proportion of mutations assigned to them), the proportion of samples in which they have detectable activity, and how tissue-specific their activity profiles were.

All six DAMUTA Misrepair signatures were active above the 0.05 activity threshold in at least 50% of samples in 19 of the 23 tissue types (Fig. 6a, Supplementary Figs. S11, S14), with a few exceptions. These exceptions – skin, lung, and esophageal – were cancer types with characteristic mutational processes that contributed the vast majority of their mutations. In skin cancer, signatures M6 accounted for the majority (>50%) of Misrepair activity in terms of number of mutations in 519/658 (79%) of samples. This signature has similar substitution profiles from UV damage-associated COSMIC SBS signatures SBS7a and SBS7b. In lung, signatures M4 and M5 together accounted for the majority (>50%) of mutations which we attributed to APOBEC activity and tobacco smoke exposure, respectively. In esophagus, signature M2, attributed to repair of reactive oxygen species (ROS), was most often the highest activity signature (174/236, 74% of samples) and accounted for the majority of mutations (>50%) in 58/236 (25%) of samples.

**Figure 6.**
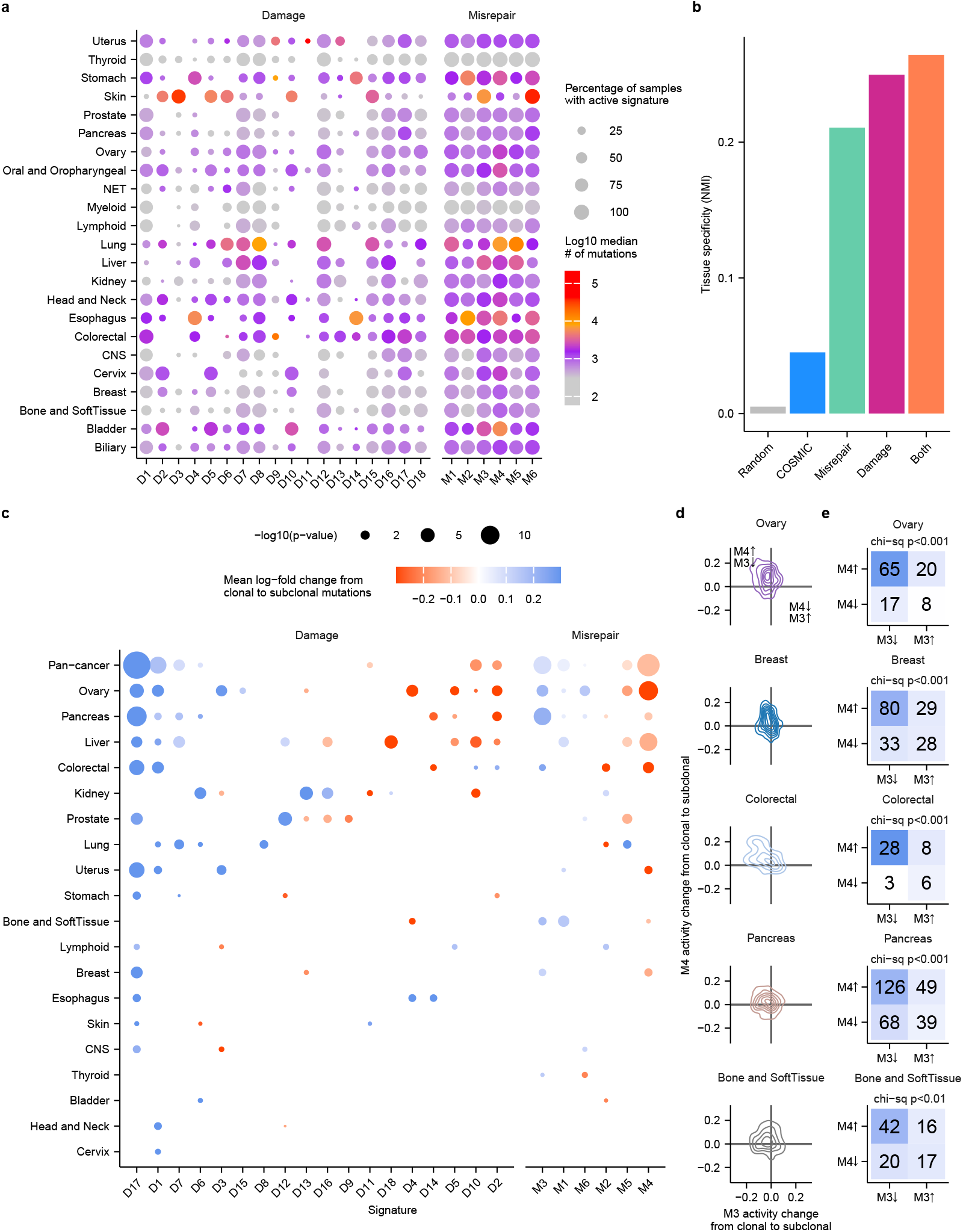
Tissue specificity and activity changes over subclonal evolution. **a**, Distribution of signature activities across tissues. Colour indicates median number of mutations attributed to each signature per tissue. Size of circle indicates proportion of samples with signature activity ≥ 0.05. **b**, Normalized mutual information of organ labels and clusters derived from hierarchical clustering. “Both” refers to the concatenation of Damage and Misrepair features. **c**, Bubble plot summarizing the mean log-fold change in signature activity over subclonal evolution for each organ. P-values are Benjamini-Hochberg corrected and only those below 0.05 are visualized. **d**, Absolute change in activity of M3 and M4 for five organ types with significant mean change in these signatures. **e**, Contingency tables for number of samples and direction of change in M3 and M4 activity in five organ types. Chi-sq P-values are Benjamini-Hochberg corrected.

In contrast, Damage signature activities were more variable across tissue types. For example, signature D11 (POLE-mutant-like) was only active in uterine, oral and oropharyngeal, head and neck, and colorectal cancers, with high mutation counts in a subset of uterine cancers. In most (17/23) types, seven or fewer damage signatures were active in more than 50% of samples. We noted select Damage signatures with the same aetiology which were active together across tissues: APOBEC signatures D2/5/10, ROS signatures D4/14. Strikingly, there was no Misrepair/Damage combination that varied identically across all tissues, underscoring the value of modelling them separately.

These activity patterns suggest that DNA damage processes have greater tissue specificity than misrepair mechanisms. To directly assess the the overall tissue specificity of Damage and Misrepair profiles, we clustered samples by their activities into 23 groups and computed the normalized mutual information (NMI) with the 23 organ type labels. Overall, Damage signature activity was more tissue-specific than Misrepair, and additionally concatenating both the Damage and Misrepair activities slightly increased NMI (Fig. 6b). Although COSMIC signature activities had higher tissue-specificity than a random labelling, they were less tissue-specific than DAMUTA’s Damage, Misrepair or Damage+Misrepair activity profiles.

### Damage and Misrepair signature activity dynamics over subclonal evolution

We next sought to examine the temporal dynamics of Damage and Misrepair signatures during cancer evolution. We ran TrackSigFreq ^15^ on 2,778 PCAWG samples to identify groups of clonal and subclonal mutations based on their variant allele frequency and, potentially, COSMIC SBS signature activities. We identified 2,014 samples with more than one clone, and from each sample analysed the cluster of mutations with the highest VAF, which would constitute the clonal mutations, and the cluster of subclonal mutations with the lowest VAF.

We then computed the Damage and Misrepair activities separately on each clonal and subclonal cluster, and identified clonal-subclonal activity changes across all samples and for each organ independently using a multinomial-regression based approach ^6^ (Fig. 6c). We found that 7/18 Damage signatures and 4/6 misrepair signatures significantly changed in activity over cancer evolution in the pan-cancer cohort, and every signature had a significant activity change in at least one organ type (P<0.05, Benjamini–Hochberg adjusted robust score test). Notable changes included a pan-cancer decrease in activity of clock-like D17, a decrease in MMRD-like D1, and an increase in APOBEC-like D2, D5, D10 in select tissues with APOBEC activity. We also noted a pan-cancer decrease in M3 and increase in M4 over subclonal evolution.

The decrease in M3 and increase in M4 was consistent across all organ types with a change in these signatures, and so we investigated the extent to which these signatures had an inverse relationship over subclonal evolution. For each of the 5 organ types with both a significant decrease in M3 and increase in M4, we asked how often these changes in activity co-occurred in the same sample. We found that while the absolute activity change did not appear to be tightly coupled (Fig. 6d), these events did frequently co-occur in the same sample (Fig. 6e, all P<0.01, chi-square test, Benjamini–Hochberg correction)

We observed organ-to-organ variability in the number of signatures with activity changes for both Damage (0–9) and Misrepair (0–5). In kidney cancers, 7 Damage signatures and only 1 Misrepair signature changed. On the other hand, 0 Damage signatures and 2 Misrepair signatures changed in Thyroid cancers. Oral and Oropharyngeal, NET, Myeloid, and Biliary did not exhibit significant changes in Damage, or Misrepair signature activities. These observations indicate that mutational process changes over tumour evolution may be emphasized in either Damage or Misrepair signatures, and that the balance between the two can be organ-specific.

## Discussion

This study presents DAMUTA, a novel probabilistic framework that for the first time successfully separates mutational signatures to represent the effects of DNA damage processes from DNA repair deficiencies. Here we have shown that by separately modelling the damage context of a mutation and the resulting substitution, we can precisely delineate damage processes from defective or inefficient DNA damage repair. Our reanalysis of APOBEC3 and UNG perturbation experiments isolated changes to the damage processes, caused by APOBEC3A knock-outs from differences in UNG activity reflected in changes in the activity of Misrepair signatures and their connectivity with APOBEC-associated damage signatures (Fig. 3e,g). This separation is not apparent from the COSMIC SBS signature activities (Supplementary Fig. S8). More broadly, we showed that DAMUTA signature activities better predicted mutations in DDR pathways in patient samples than did COSMIC activities. Clinically, DDR mutations are significantly associated with survival ^38^. Importantly, Misrepair signatures were more informative than Damage signatures in determining DDR-deficient cancers, and together our results demonstrate the effectiveness of DAMUTA for partitioning subjects based on DNA damage or misrepair processes.

Our reanalysis of SBS mutation spectra from 18,947 tumours from Genomics England, PCAWG, and Hartwig Foundation datasets revealed some expected and some surprising patterns of DNA damage and the cellular response in tumours and during their development. Some SBS signature activities are highly tissue-specific ^1^, and our analysis revealed that this tissue-specificity arises primarily from differences in damage exposure rather than repair, as one would expect if DNA damage response is relatively consistent across tissue types. However, our analysis suggests some consistent sample-specificity in DDR. For example, among our 30 connectivity patterns there are three that match SBS1. These patterns have a shared Damage context, which is highly enriched for the CpG dinucleotides, but strong preference for distinct Misrepairs.

The previously reported widespread tissue-specificity of SBS signatures may be due to subtle variations in the activity of repair pathways, or the interactions between damage and repair. These variations are naturally captured in DAMUTA whereas the COSMIC signatures, like our connection patterns, are capturing major modes of interaction between damage and misrepair. Surprisingly, we found no clear surrogate among our connection patterns for SBS5, a SBS signature with unknown aetiology which is ubiquitously active in tumors. The most similar connection patterns link to the transition Misrepair signature (M3), but there is no consistent Damage signatures among the closest matches. More broadly, only some SBS signatures had clear surrogates in connection patterns (Fig. 5), and this proportion does not increase when we add more connection patterns (Supplementary Fig. S12). This points to clear distinctions in how DAMUTA is capturing shared patterns of SBS mutations.

Many SBS signatures vary in their activity throughout tumour evolution, though not always in a consistent direction ^9^. In contrast, we found much more consistent changes in the activities of select Misrepair and Damage signatures. Interestingly, we find that M3, which consists solely of transitions, dominates early in tumour development whereas M4, a near-uniform pattern of substitutions, dominates late. One possibility is that M4 is capturing translesion synthesis by Y-family polymerases, and to support more rapid proliferation, cancer cells rely increasingly on this mechanism to resolve lesions during replication. This tendency has been previously proposed in C. elegans ^35^, and might also explain M4’s increased activity in the UNG high samples; as suggested by Petljak *et al.* ^23^, efficient uracil removal may lead to more frequent replication across abasic sites (Supplementary Fig. S7).

DAMUTA represents the sequence context preferences of Misrepair signatures through interaction with Damage signatures. This makes DAMUTA’s signatures more identifiable than previous approaches ^37^, however it also means DAMUTA does not directly capture Misrepair context. We see several opportunities to extend Misrepair signatures to directly consider contextual features: first, adjacent nucleotide sequence may be included which would require explicitly deconvolving these compound effects with DNA damage; second, modifying the generative process to consider additional genomic features such as transcription- or replication-strandedness, replication timing, nucleosome occupancy, or DNA conformation; last, incorporation of additional data types, such as those generated by recent investigations of DNA lesions which persist through multiple rounds of replication ^2,3^. We expect these enhancements will support the development of new models, with more biologically-informed inductive biases.

By providing a more interpretable representation of the mutation processes in tumours, DAMUTA paves the way for a more unified and mechanistic understanding of cancer mutational signatures, as well as a more succinct summary of these processes. We propose DAMUTA as a replacement or supplement for the COSMIC SBS signature framework, and anticipate it will advance the dissection of molecular etiologies of mutation signatures and the identification of patients with DNA repair deficient cancers for synthetic lethality-based therapy. To aid in its wide adoption, we provide freely-available open source code to fit signatures to different datasets, as well as an annotated set of damage and misrepair signatures on our tumour set and a set of common interaction matrix patterns.

## Acknowledgment

KRC reports consulting fees received from Abbvie Inc. and research funding from Sanofi and Standard BioTools, all unrelated to this work. QM is supported by NIH/NCI Cancer Center Support Grant P30 CA008748 (Vickers). CFH is supported by the ACM SIGHPC Computational and Data Science Fellowship, NSERC doctoral fellowship, and the Data Sciences Institute at the University of Toronto.

## Data and code availability

Data and code required to reproduce this study can be retrieved from: https://github.com/morrislab/damuta

## Methods

### Dataset curation

We estimated DAMUTA signatures and activities for 18,974 whole genome sequenced tumours originating from 23 organs (Table S1) with a median of 6878 mutations per sample (Figure S1). Making use of mutation catalogues compiled by Degasperi et al. ^8^ we sourced mutation counts from four datasets: Pan Cancer Analysis of Whole Genomes (PCAWG) ^32^, Hartwig Medical Foundation (HMF) ^25^, Genomics England 100,000 Genomes Project (GEL) ^33^ and International Cancer Genome Consortium (ICGC) ^16^. These datasets represent samples from a variety of histologies, pre- and post-treatment samples, and samples from primary and metastatic tumours. These datasets consist predominantly of patients of white European ancestry, and male and female patients are included.

### Calculation of marginalized profiles

A marginalized profile of a size-96 SBS signature is calculated by summation. For a given SBS signature S, the substitution profile was calculated by summing mutation probabilities across all trinucleotide contexts for each of the six possible pyrimidine substitutions (C>A, C>G, C>T, T>A, T>C, T>G):

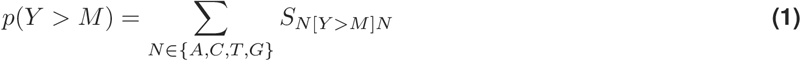

where *Y* is the reference pyrimidine (C or T) and M is the mutated base.

The trinucleotide context profile was calculated by summing mutation probabilities across all possible substitutions for each trinucleotide context (AAA, ACA, … TTT):

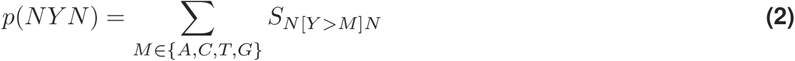

where NYN represents one of the 32 possible trinucleotide contexts centered on the reference pyrimidine Y and M is the mutated base.

### Determining C/T bias

We call a damage signature as C-context biased if the ratio of total weight is C:T > 2, and T-biased if <0.5. Due to the constraints of the DAMUTA model, all misrepair signatures must have both C and T elements appearing together, as each type sums to 1 (Fig. 2b).

### Estimation of COSMIC activities using DeconstructSigs

DeconstructSigs v1.9.0 was run on mutation counts for 18,947 tumours with COSMIC v3.2 for GRCh37 provided as reference signatures (n=78), with parameter settings `contexts.needed=T and signature.cutoff=0.06`which is the default cuttoff threshold.

### Estimation of Damage and Misrepair activities using DAMUTA

DAMUTA models per-genome damage probabilities as *θ* and per-genome, per-damage misrepair probabilities are captured by the rows of the *A* matrix which fits the relationships between Damage and Misrepair signatures. For each mutation, depending on the assigned Damage signature, DAMUTA uses the appropriate Misrepair probability distribution. We use the product of these, the interaction matrix (*θA*), to calculate overall *activity* of Misrepair signatures and analyze the individual behaviours of these two signature types. We define DAMUTA signature activities as:

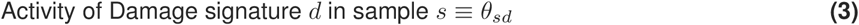

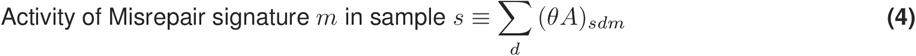

### Generative process

The generative process for DAMUTA is as follows: for each genome, *g*, DAMUTA assigns activities to the damage processes, *θ*_*g*_, and damage-misrepair association matrix *A*_*g*_, which specifies for each damage process the relative proportion of its damage that are misrepaired by the different misrepair signatures. Latent factors are individually described in Table S2.

1. Define damage signatures by drawing *ϕ*_*d*_ *∼ Dirichlet*(*π*) ∀*d* ∈ {1, …, *D*}
2. Define misrepair signatures by drawing *η*_*m*_ *∼ Dirichlet*(*λ*) ∀*m* ∈ {1, …, *M*}
3. For each tissue-type *t* ∈ {1, …, *T*}:
  a. Draw the tissue-specific prior on misrepair signature activity according to *a*_*t*_ *∼ Gamma*(*α*_*a*_, *β*_*a*_) and *b*_*t*_ *∼ Gamma*(*α*_*b*_, *β*_*b*_)
  b. For each sample *s* ∈ {1, …, *S*} of type *t*:

i. Draw the Tissue-specific association prior Γ_*s*_ *∼ Gamma*(*a*_*t*_, *b*_*t*_)
ii. Draw the Damage signature activities *θ*_*s*_ *∼ Dirichlet*(*ψ*)
iii. Draw the Damage/Misrepair signature associations *A*_*s,d*_ *∼ Dirichlet*(Γ_*s*_)
iv. Draw the mutation type counts *Y*_*s*_ *∼ Multinomial*(*P*_*s*_, *N*_*s*_) where *P*_*s*_ = concat(*θ*_*s*_*ϕ*^(*C*)^ ⊗ *θ*_*s*_*A*_*s*_*η*^(*C*)^, *θ*_*s*_*ϕ*^(*T*)^ ⊗ *θ*_*s*_*A*_*s*_*η*^(*T*)^)

In the de-novo setting, signature definitions are jointly learned by meanfield automatic differentiation variational inference (ADVI) ^18^ at the same time as local parameters (*e.g.* activities). Signatures can be initialized uniformly at random, or by starting from k-means cluster centroids found from the data. A k-means initialization converges to a solution faster than random initialization, and finds more stable Damage Signatures (Fig. S5). It is also possible to initialize DAMUTA from the marginalized context and substitution profiles of known signatures (For example, COSMIC signatures as shown in Fig. 1c).

In the fixed-signature setting, user-provided *ϕ* and *η* are passed as “observed” parameters to pymc3^29^, which constrains the signature definitions to those provided, and fits only the activities.

To fit DAMUTA to 18,947 samples from PCAWG, Hartwig (HMF), and Genomics England (GEL), we use the following parameter settings:

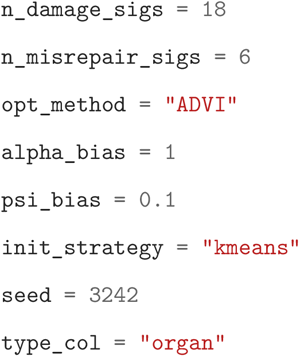

### Selecting number of signatures

We used BIC to determine an appropriate setting for the number of signatures (*D* and *M* in the plate model, Figure 1b) and selected the smallest model within 0.95% log likelihood of the best fitting model. The number of signatures selected was also supported by examining elbow-curves of hierarchically clustered marginalized COSMIC signatures (Figure S2) and by examining how likelihood plateaus with increasing *D* and *M* on average over a hyperparameter sweep (Fig. S4).

In settings where it is known a-priori which COSMIC signatures are present in a sample, it may be preferable to cluster these profiles and initialize DAMUTA using these. This approach is also recommended when fitting DAMUTA de-novo in settings where all samples have fewer than 1000 mutations.

We threshold low activities in a similar manner as DeconstructSigs ^27^. Thresholds were chosen to balance i) maximizing the total activity per sample ii) minimizing the number of active signatures per sample. For the Damage and Misrepair activities we take a threshold of 0.05. We do not threshold activities in interaction matrices, which tend to contain many low-value entries: 0.05 represents the 97.6th percentile of these values (Fig. S14).

### Tissue specificity of signature activity

To quantify the tissue specificity of damage and misrepair, we performed hierarchical clustering with cosine distance and complete linkage on signature activities from DAMUTA and COSMIC. We cut the cluster dendrogram to produce clusters according to the number of organs in the data (n=23), and computed normalized mutual information using the sample organ labels (Fig. 6b).

### RadEmu multinomial regression on subclonal activity changes

For each organ, RadEmu v2.0^6^ was fit to test the association of activity changes from first-to-last subclone. Robust test score was used to estimate P-values.

### NMF extraction of 30 connection patterns

We picked 30 NMF factors by examining an elbow plot of reconstruction error, and signature stability as measured by average silhouette score over 3 seeds. The number of COSMIC signatures which could be matched with >0.8 cosine similarity to a connection pattern also appeared to plateau after approximately 30 (Fig. S12).

We fit connection patterns using Elastic net regularized NMF, with parameters `alpha_w=1e-6, alpha_h=5e-6, solver=“cd”`. These settings were picked to maximize total activity per sample, while minimizing the number of active signatures in each sample. Connection patterns and activities were then normalized to sum to 1.

### Classification of repair-pathway altered samples

To calculate signature mutation counts for DAMUTA and COSMIC feature sets, we estimated activities using DAMUTA, and DeconstructSigs respectively, and scaled these activities by the number of mutations in each sample. The resulting values are not rounded to integer values. This resulted in 24 DAMUTA (18 Damage, 6 Misrepair) and 78 COSMIC-based features.

Gradient boosted machines were trained using LightGBM v3.3.4 with n_estimators=30 and all other parameters set to defaults. Each classifier was re-trained 10 times in parallel, with 30% of the samples withheld for model evaluation. Data were randomly stratified on the positive class to ensure examples were evenly split across train and test sets.

