## supplemental figures and tables for "Damage and Misrepair Signatures: Compact Representations of Pan-cancer Mutational Processes"

### Supplementary Figures

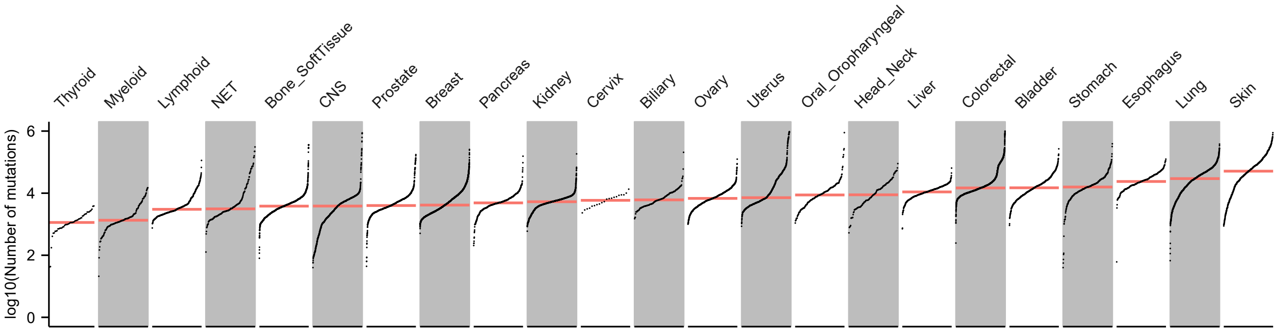


**Supplementary Figure S1.** Number of mutations per sample. Red line indicates median number of mutations for each organ.

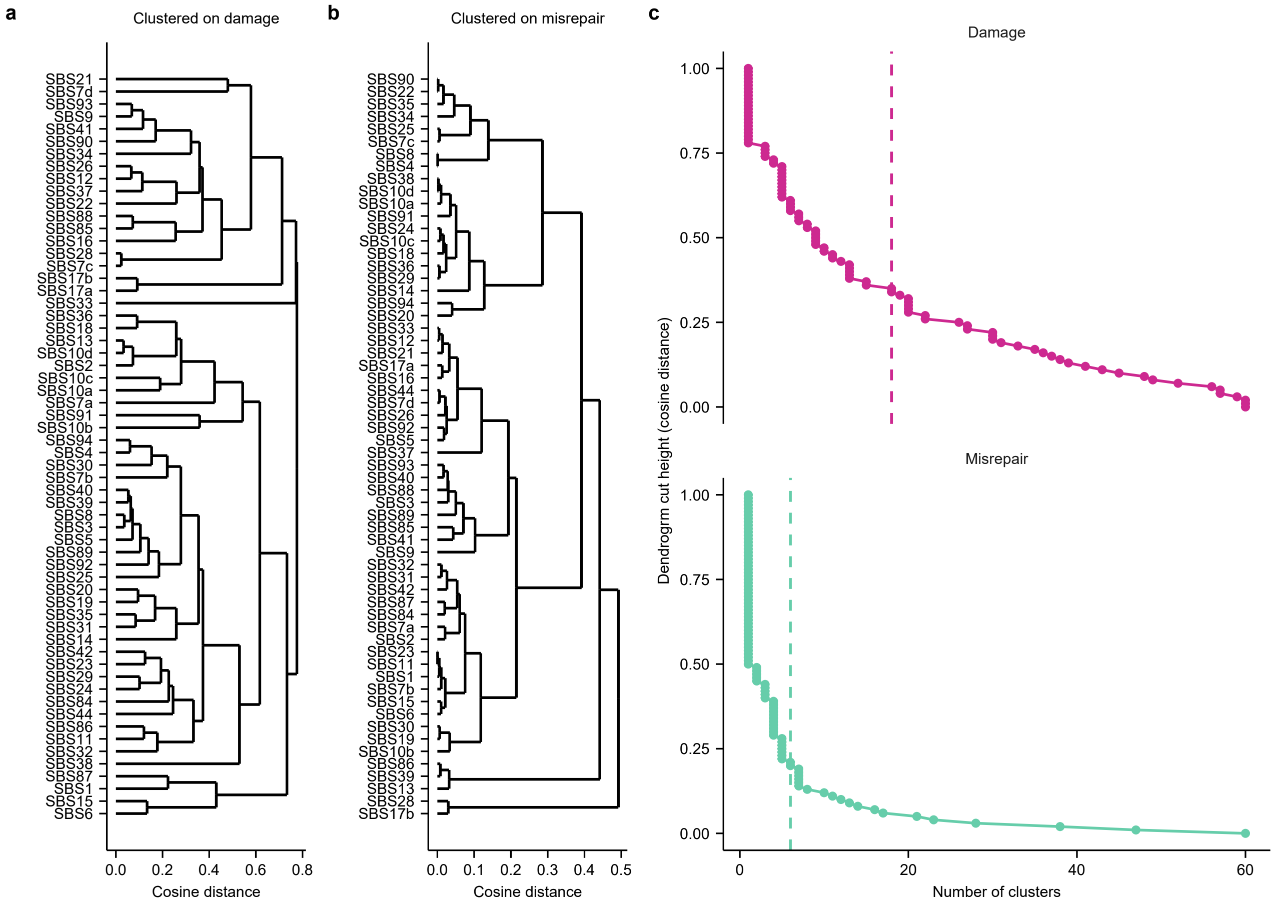


**Supplementary Figure S2.** **a**, COSMIC signatures clustered by their trinucleotide context profiles using average linkage and cosine distance. **b**, COSMIC signatures clustered by their substitution profiles using average linkage and cosine distance. **c**, Elbow plots of number of clusters defined by dendrogram cut. Dashed vertical lines indicate choice of number of signatures used in denovo extraction: 18 Damage, 6 Misrepair.

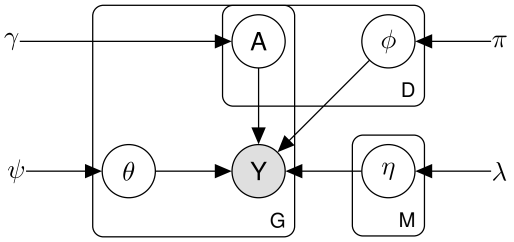


**Supplementary Figure S3.** DAMUTA graphical model with uninformative prior on A.

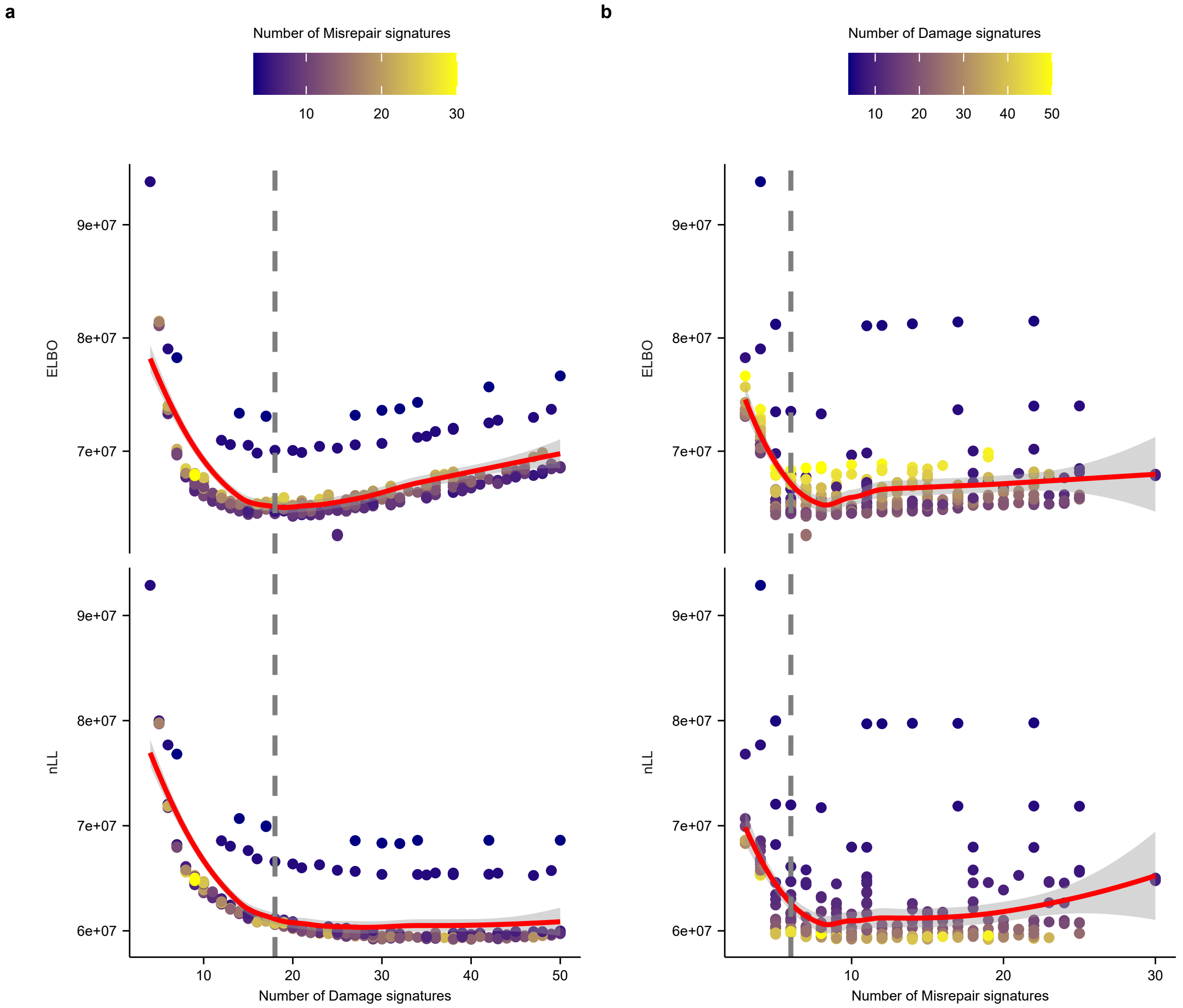


**Supplementary Figure S4.** Evidence lower bound (ELBO, top) and negative log likelihood (nLL, bottom) for a hyperparameter sweep over choice of $D$ and $M$. Red solid line represents a LOESS-smoothed fit to the data. **a** Dashed grey line indicates the selected number of Damage signatures (18). **b**, Dashed grey line indicates the selected number of Misrepair signatures (6)

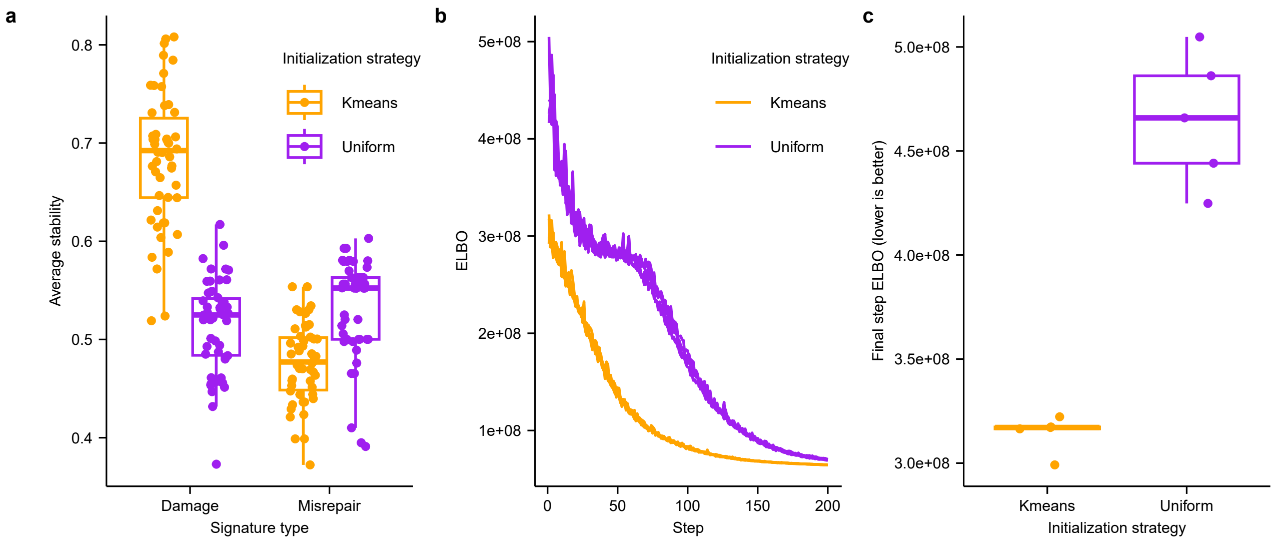


**Supplementary Figure S5.** **a**, Signature stability over 5 initialization seeds, calculated as average silhouette width on 50 rounds of kmeans clustering estimated signature definitions. **b**, ELBO curves for 5 initialization seeds. **c**, Final value of ELBO of fitted model, with 5 initializations.

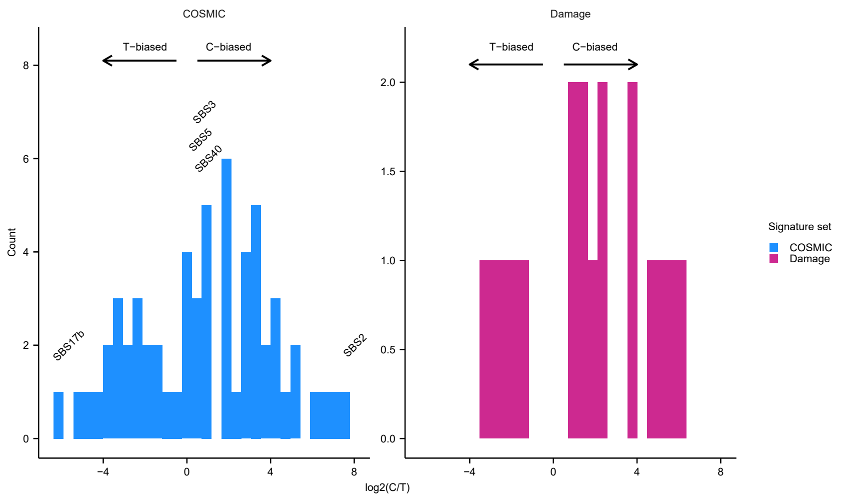


**Supplementary Figure S6.** Distribution of C/T context balances in signature sets. Left: COSMIC, with select signatures labelled. Right: DAMUTA Damage signature set

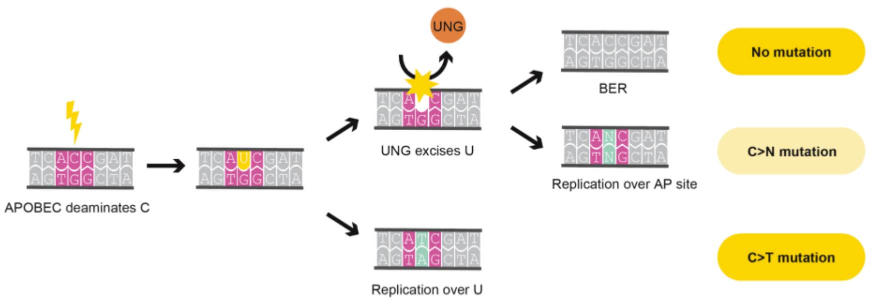


**Supplementary Figure S7.** DNA damage and repair process involving APOBEC and UNG. UNG is able to remove uracil (U) in DNA, resulting in base excision repair (BER), or replication across an abasic (AP) site. In the absence of UNG, uracil remains and is replicated across as if it is thymine.

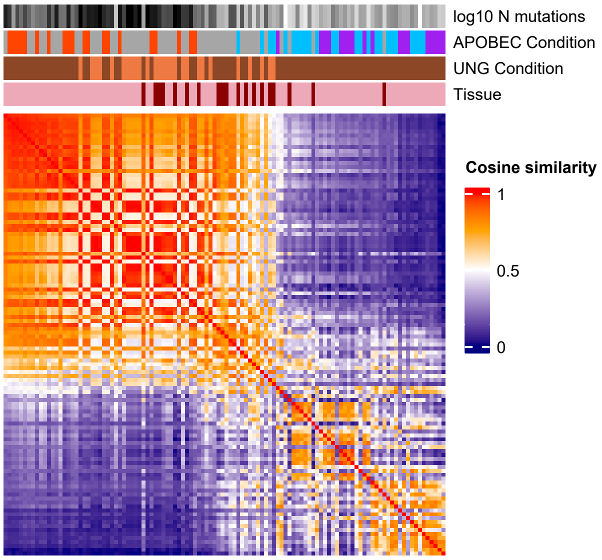


**Supplementary Figure S8.** Cosine similarity of COSMIC activities for 112 cell line samples with knockout of APOBEC or modulation of UNG. Annotation track colors as in Fig. 3a.

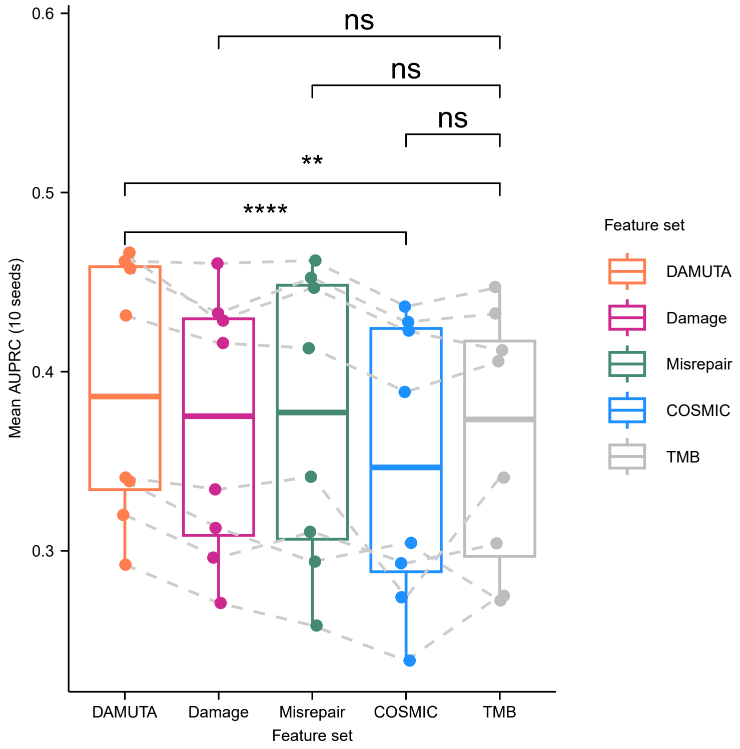


**Supplementary Figure S9.** Mean AUPRC for classifying samples across eight repair pathways, P-values for paired t-test shown: *: P<0.05, **: P < 0.01, ***: P < 0.001, ****: P < 0.0001 . DAMUTA: Damage and Misrepair activities are concatenated. TMB: tumor mutational burden.

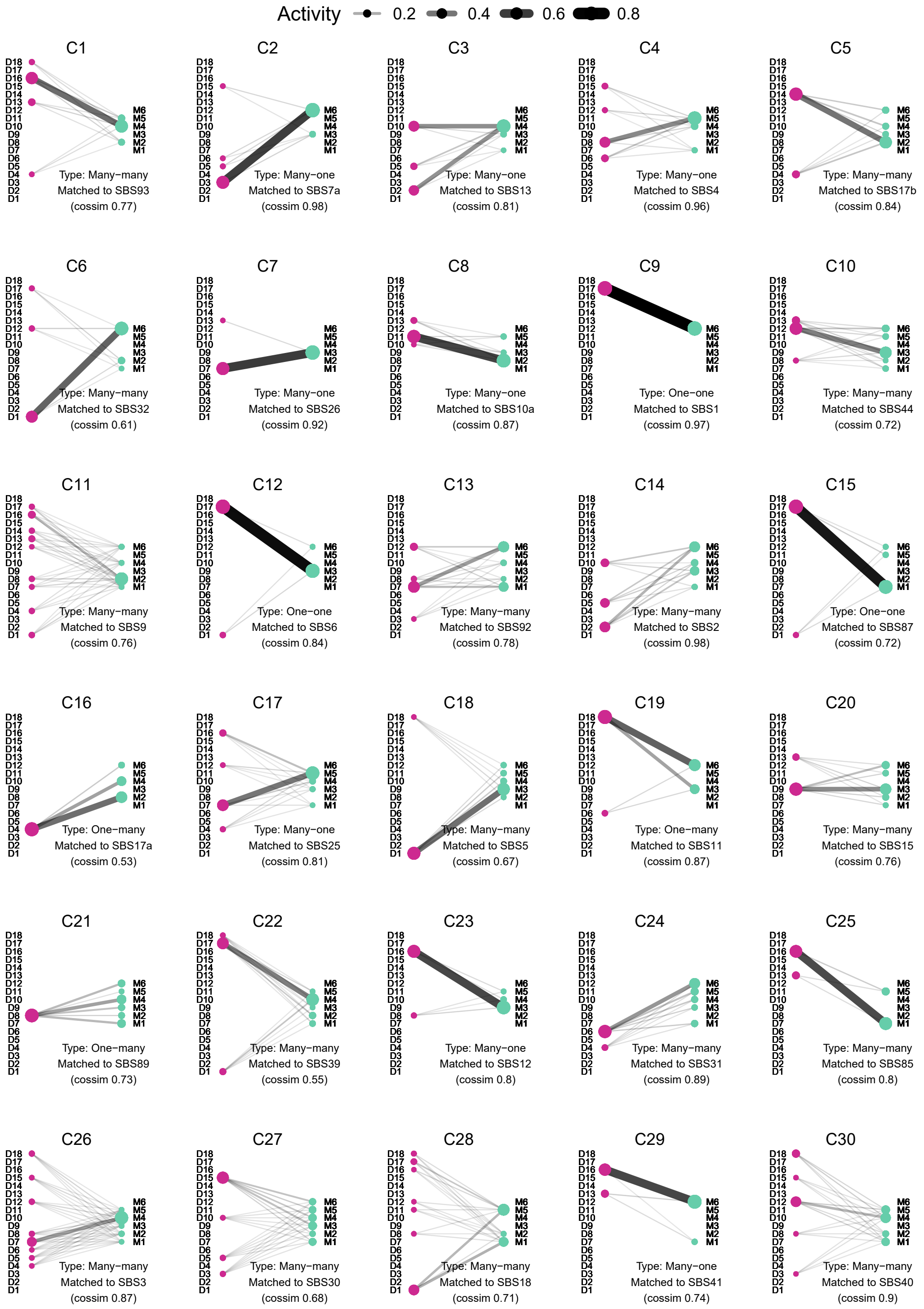


**Supplementary Figure S10.** Connection patterns extracted using NMF on DAMUTA interaction matrices (N=30). Patterns are labelled by Connection pattern name, connection type, hungarian-algoritm matched COSMIC signature, and its cosine similarity based on the Connection pattern's collapsed SBS distribution. Only connections between Damage or Misrepair signatures with >0.05 activity are visualized.

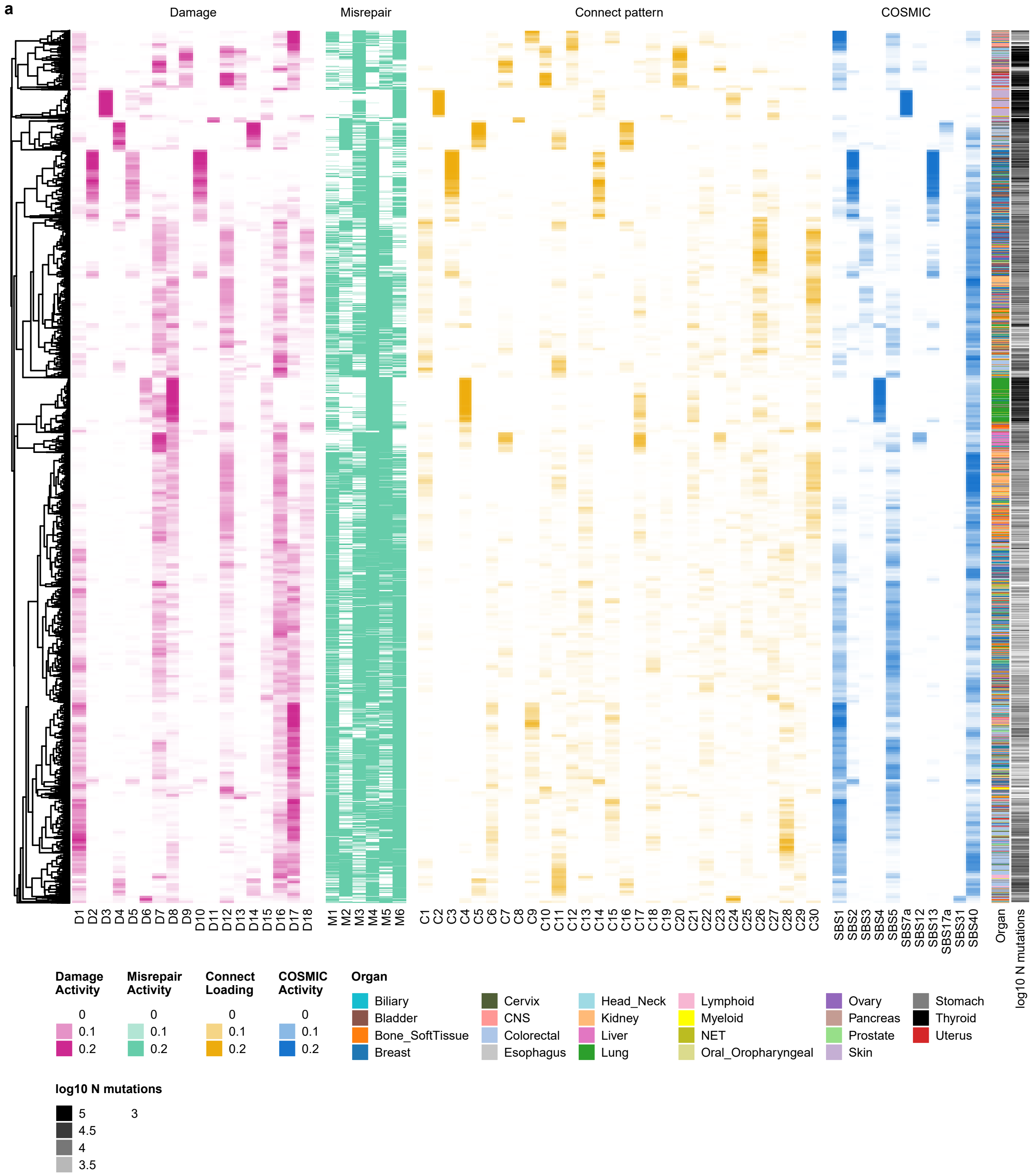


**Supplementary Figure S11.** Panorama of activities of Damage, Misrepair, Connection, and select COSMIC signatures in 18,947 tumours

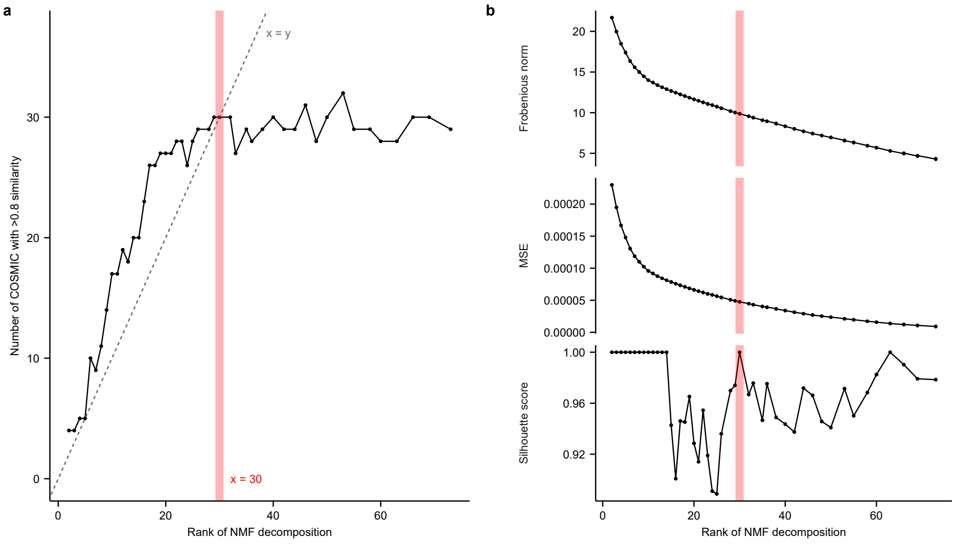


**Supplementary Figure S12.** **a**, Number of COSMIC signatures with high similarity to NMF-derived Connection signatures at different ranks for NMF decomposition. Red line indicates the selected rank for DAMUTA Connection signatures (x=30). **b**, Other metrics considered when picking number of NMF factors for connection patterns

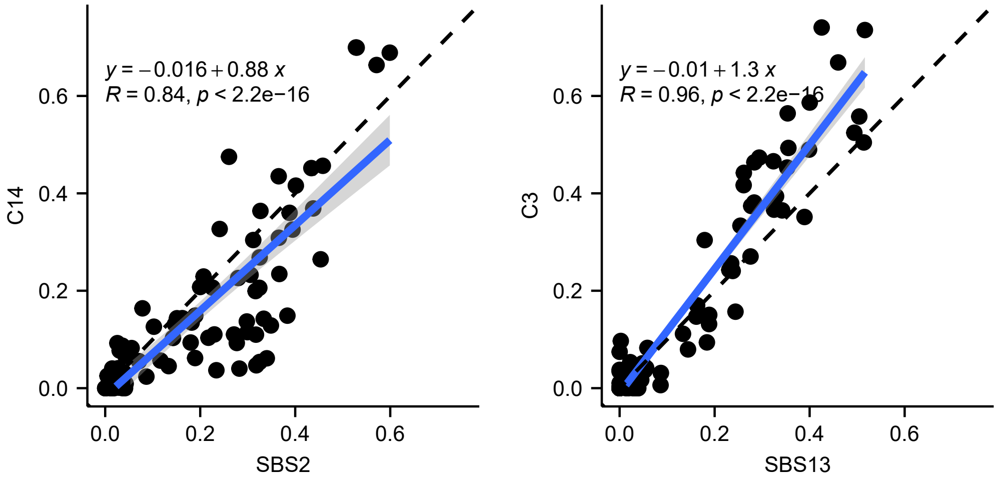


**Supplementary Figure S13.** Correlation of activity of APOBEC signatures in cell lines between **a**, Connection pattern C14 and SBS2, and **b**, Connection pattern C3 and SBS13.

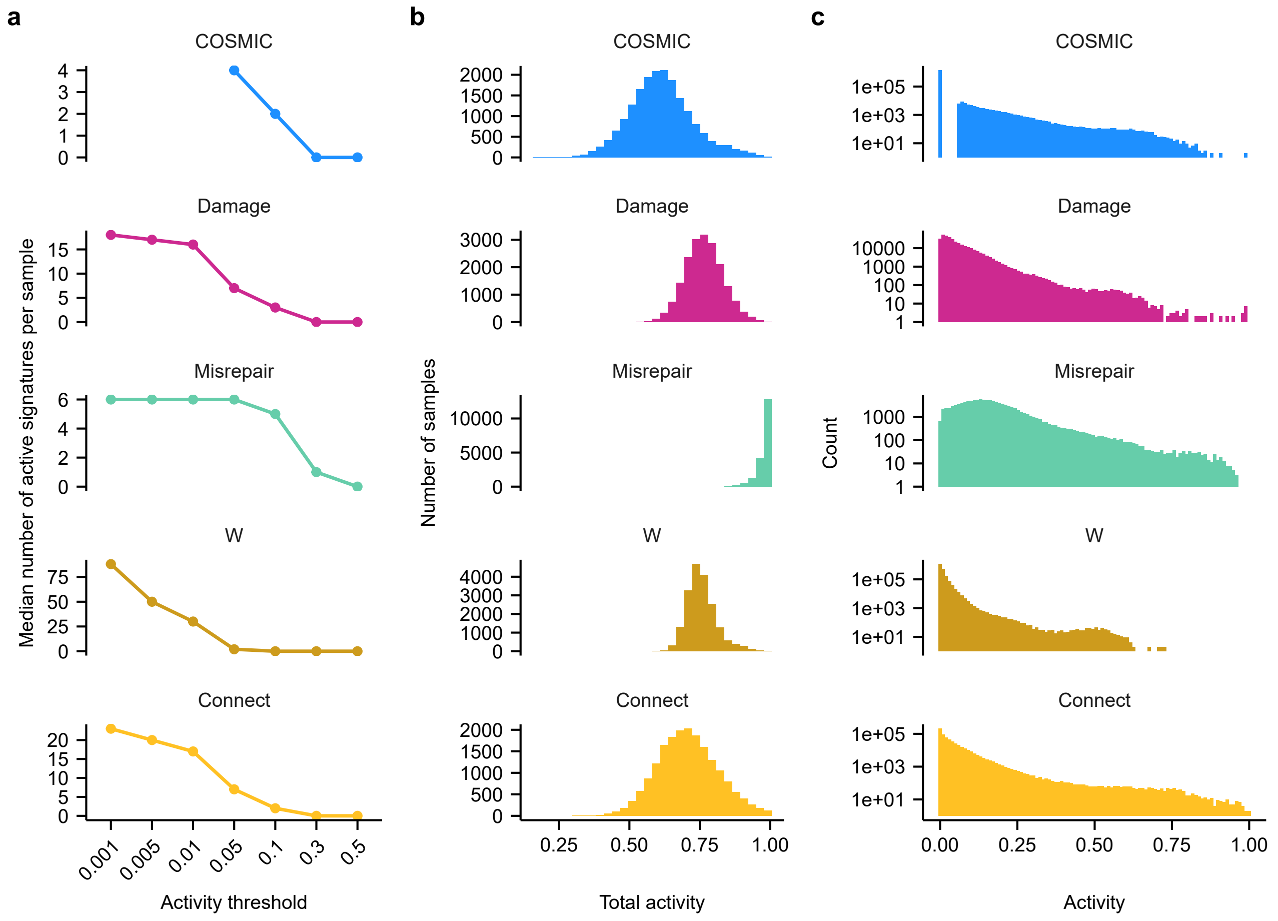


**Supplementary Figure S14.** **a**, Median number of active signatures per sample for varying choices of activity threshold. **b**, Distribution of total activities for various signature types for the chosen thresholds: COSMIC=0.06 (DeconstructSigs default), Damage=0.05, Misrepair=0.05, W=0.01, Connect=0.05. **c**, Distribution of activities for all signatures in all samples.

#### Supplementary tables

| Organ | GEL | HMF | PCAWG | ICGC | Total |
|---|---|---|---|---|---|
| Breast | 2572 | 661 | 214 | 530 | 3977 |
| Colorectal | 2348 | 493 | 60 | 0 | 2901 |
| Bone_SoftTissue | 1480 | 164 | 98 | 0 | 1742 |
| Kidney | 1355 | 100 | 189 | 0 | 1644 |
| Lung | 1009 | 368 | 86 | 0 | 1463 |
| Prostate | 311 | 368 | 286 | 0 | 965 |
| Uterus | 713 | 66 | 51 | 0 | 830 |
| CNS | 444 | 78 | 294 | 0 | 816 |
| Ovary | 523 | 144 | 113 | 0 | 780 |
| Skin | 258 | 293 | 107 | 0 | 658 |
| Bladder | 349 | 140 | 23 | 0 | 512 |
| Pancreas | 59 | 75 | 326 | 0 | 460 |
| Liver | 21 | 48 | 326 | 0 | 395 |
| Lymphoid | 181 | 0 | 202 | 0 | 383 |
| Stomach | 204 | 39 | 75 | 0 | 318 |
| Esophagus | 0 | 138 | 98 | 0 | 236 |
| NET | 92 | 100 | 0 | 0 | 192 |
| Oral_Oropharyngeal | 186 | 0 | 0 | 0 | 186 |
| Myeloid | 91 | 0 | 70 | 0 | 161 |
| Biliary | 26 | 81 | 35 | 0 | 142 |
| Head_Neck | 0 | 61 | 57 | 0 | 118 |
| Thyroid | 0 | 0 | 48 | 0 | 48 |
| Cervix | 0 | 0 | 20 | 0 | 20 |
| Total | | | | | 18947 |

**Table S1.** Organ type counts per dataset

| | Variable | Type | Meaning |
|---|---|---|---|
| Constants | G | Integer | Number of genomes |
| | T | Integer | Number of tissue types |
| | $N_{g=1,\dots,G}$ | Integer | number of mutations in genome $g$ |
| | D | Integer | Number of damage signatures |
| | M | Integer | Number of misrepair signatures |
| Parameters | $\psi_{c=1,\dots,32}$ | Positive real | prior weight of context type $c$ in a signature, 0.5 gives generally good sparsity |
| | $\lambda'_{b\in A,C,T,G}$ | Positive real | prior weight of mismatch type $b$ in a signature, 1 gives generally good uniformity |
| | $\lambda_{e=1,\dots,6}$ | Positive real | Set using $\lambda'$ such that C and T mismatches have a shared prior: $\lambda_1 = \lambda_4 = \lambda'_A; \lambda_2 = \lambda_6 = \lambda'_G; \lambda_3 = \lambda'_T; \lambda_5 = \lambda'_C$ |
| | $\psi_{d=1,\dots,D}$ | Positive real | prior weight of damage signature $d$ in a sample, 0.1 gives generally good sparsity |
| | $\alpha_a$ | Positive real | shape prior on shape parameter for tissue-type specificity of misrepair signature activity |
| | $\beta_a$ | Positive real | rate prior on on shape parameter for tissue-type specificity of misrepair signature activity |
| | $\alpha_b$ | Positive real | shape prior on on rate parameter for tissue-type specificity of misrepair signature activity |
| | $\beta_b$ | Positive real | rate prior on on rate parameter for tissue-type specificity of misrepair signature activity |
| Factors | $\phi_{d=1,\dots,D,c=1,\dots,32}$ | Probability on [0,1] | probability of context type $c$ appearing in damage signature $d$ |
| | $\eta_{m=1,\dots,M,e=1,\dots,6}$ | Probability on [0,1] | probability of mismatch type $e$ appearing in misrepair signature $m$ |
| | $A_{s=1,\dots,S,d=1,\dots,D,m=1,\dots,M}$ | Probability on [0,1] | probability (activity) of the $m$-th misrepair signature acting on the $d$-th damage signature, in sample $s$ |
| | $\theta_{s=1,\dots,S,d=1,\dots,D}$ | Probability on [0,1] | probability (activity) of the $d$-th damage signature occurring in sample $s$ |
| | $\Gamma_{s=1,\dots,S,m=1,\dots,M}$ | Positive real | relative weight (transformed activity) of the $m$-th misrepair signature occurring in sample $s$ |
| | $a_{t=1,\dots,T}$ | Positive real | shape parameter for tissue-type specificity of misrepair signature activity |
| | $b_{t=1,\dots,T}$ | Positive real | rate parameter for tissue-type specificity of misrepair signature activity |
| | $Y_{s=1,\dots,S,i=1,\dots,96}$ | Integer | count of variants of the $i$-th mutation type (as a function of the $c$-th context type and $e$-th mismatch type), in sample $s$ |

**Table S2.** Description of factors in graphical model
